# Unsupervised decoding of single-trial EEG reveals unique states of functional brain connectivity that drive rapid speech categorization decisions

**DOI:** 10.1101/686048

**Authors:** Rakib Al-Fahad, Mohammed Yeasin, Gavin M. Bidelman

**Affiliations:** Department of Electrical and Computer Engineering, University of Memphis, Memphis, 38152 TN, USA; Institute for Intelligent Systems, University of Memphis, Memphis, TN, USA; School of Communication Sciences & Disorders, University of Memphis, Memphis, TN, USA; University of Tennessee Health Sciences Center, Department of Anatomy and Neurobiology, Memphis, TN, USA

**Keywords:** Categorical speech perception, machine learning, speech processing, stability selection, functional connectivity

## Abstract

Categorical perception (CP) is an inherent property of speech perception. The response time (RT) of listeners’ perceptual speech identification are highly sensitive to individual differences. While the neural correlates of CP have been well studied in terms of the regional contributions of the brain to behavior, functional connectivity patterns that signify individual differences in listeners’ speed (RT) for speech categorization is less clear. To address these questions, we applied several computational approaches to the EEG including graph mining, machine learning (i.e., support vector machine), and stability selection to investigate the unique brain states (functional neural connectivity) that predict the speed of listeners’ behavioral decisions. We infer that (i) the listeners’ perceptual speed is directly related to dynamic variations in their brain connectomics, (ii) global network assortativity and efficiency distinguished fast, medium, and slow RT, (iii) the functional network underlying speeded decisions increases in negative assortativity (i.e., became disassortative) for slower RTs, (iv) slower categorical speech decisions cause excessive use of neural resources and more aberrant information flow within the CP circuitry, (v) slower perceivers tended to utilize functional brain networks excessively (or inappropriately) whereas fast perceivers (with lower global efficiency) utilized the same neural pathways but with more restricted organization. Our results showed that neural classifiers (SVM) coupled with stability selection correctly classify behavioral RTs from functional connectivity alone with over 90% accuracy (AUC=0.9). Our results corroborate previous studies by confirming the engagement of similar temporal (STG), parietal, motor, and prefrontal regions in CP using an entirely data-driven approach.

## INTRODUCTION

When identifying speech, listeners naturally group sounds into smaller sets of discrete (phonetic) categories through the process of categorical perception (CP) (Harnad and Bureau, 1987; Liberman et al., 1967; Pisoni, 1973; Pisoni and Luce, 1987). Presumably, this type of behavioral “downsampling” promotes speech comprehension by generating perceptual constancy in the face of enormous physical variation in multiple acoustic dimensions, e.g., talker variability in tempo, pitch, or timbre (Prather et al., 2009). CP is often characterized by sharp (stair-stepped) identification and peaked (better) discrimination functions near the categorical boundary when classifying an otherwise equidistant acoustic continuum.

Germane to the present study, response time (RT) data also reveal differences in the speed of listeners’ of categorical decisions (Bidelman et al., 2013; Pisoni and Tash, 1974). In perceptual labeling tasks, for example, listeners categorize prototypical speech sounds (e.g., exemplars from their native language) much faster than their ambiguous or less familiar counterparts (e.g., nonnative speech sounds) (Bidelman and Lee, 2015). RTs also slow near perceptual speech boundaries, where listeners shift from hearing one linguistic class to another (e.g., /u/ vs. /a/ vowel) and presumably require more time to access the “correct” speech template (Bidelman et al., 2013; Liebenthal et al., 2010; Pisoni and Tash, 1974; Reetzke et al., 2018). Relatedly, RTs vary with task manipulations and individual differences in speech perception in different populations. Studies demonstrate listeners’ speed in speech identification is highly sensitive to stimulus familiarity (Bidelman and Walker, 2017; Liebenthal et al., 2010; Lively et al., 1993), auditory plasticity of short-(Liberman et al., 1967) and long-term (Bidelman et al., 2014b; Bidelman and Alain, 2015; Bidelman and Lee, 2015) experience, and neuropathologies and language-learning disorders (e.g., (Bidelman et al., 2017, 2014a; Calcus Axelle et al., 2016; Hakvoort Britt et al., 2016)). Given its fundamental role in the perceptual organization of speech, understanding individual differences in CP and its underlying neurobiology is among the broad interests to understand how sensory features are mapped to higher order perception (Bidelman et al., 2013; Phillips, 2001; Pisoni and Luce, 1987).

The neuronal elements of the brain organize in complicated structural networks (Cajal, 1995). Increasingly, it is appreciated that anatomical substrates constrain the dynamic emergence of coherent physiological activity that can span multiple spatially distinct brain regions (Bressler, 1995; Bullmore and Sporns, 2009; Fries, 2005). Such densely intra-connected, sparsely inter-connected, dynamic connected networks are thought to provide the functional basis for information processing, mental representations, and complex behaviors (Bassett and Bullmore, 2006; Honey et al., 2007; Newman, 2003; Tononi et al., 1994). In this regard, neuroimaging studies have identified several functional brain regions that are important to CP including primary auditory cortex, left inferior frontal areas (i.e., Broca’s area), and middle temporal gyri (e.g., Guenther Frank H. et al. 2004; Binder et al. 2004; Myers et al. 2009; Chang et al. 2010; Liebenthal et al. 2010; Bidelman and Lee 2015; Alho et al. 2016; Toscano et al. 2018). Previous studies also suggest that more neurons are preferentially activated by the prototypes of the speech categories compared to those at category boundaries (Guenther and Gjaja, 1996). Similarly, improved discriminability at category boundaries could reflect an increased number of neurons encoding sensory cues at these perceptual transitions (Bauer and Der, 1996; Guenther et al., 1999). Such neuronal overrepresentations warp the sensory space and may account for the aforementioned RT effects in speech categorization. Still, while the neural correlates of CP have been well studied in terms of the regional contributions to behavior, we are aware of no studies that have investigated the mechanisms of speech CP from a full-brain (functional connectivity) perspective. Here, we focus on the speed (RT) of listeners perceptual speech identification as RTs are highly sensitivity to individual differences in CP (Bidelman et al., 2014b, 2014a; Bidelman and Alain, 2015; Bidelman and Walker, 2017) and reflect an objective, continuous measure of perceptual categorization skill.

Functional connectivity matrices derived from neuroimaging data are highly sparse and reflect high dimensional data. Hence, finding RT-related network edges is challenging. State-of-the-art studies usually use naive approaches to discover and analyze each edge individually and then compensate for possible errors arising from multiple comparisons (e.g., family-wise error or false discovery rate). These studies mostly yield an unstable set of network edges that are highly sensitive to changes in the hyperparameters within and between datasets (e.g., neural responses from different populations). In this regard, variable selection attempts to identify the most salient subset of variables from a larger set of features mixed with irrelevant variables. This problem is especially challenging when the number of available data samples is smaller compared to the number of possible predictors. Using generic subsampling and high-dimensional selection algorithms, stability selection can yield a stable set of features that distinguish subgroups of the data (e.g., here, listeners with slow vs. fast perceptual decisions). It has widely been used in diverse fields of science including gene selection and neuroimaging. One of the downsides of multivariate approaches is that outcomes often depend on model parameters (e.g., regularization factor). Compared to conventional multivariate approaches, stability selection produces more reliable estimations because of its internal randomization implemented as bootstrap-based subsampling (Meinshausen and Bühlmann, 2010; Shah and Samworth, 2013). Here, we propose a systematic approach to determine and rank RT-related functional connectivity among brain regions that are consistent across model parameters. In doing so, we identify, objectively, the most important properties (i.e., features) of the functional EEG connectome that describe perceptual processing with regard to categorization.

The primary aim of this study was to test whether individual differences in speeded speech categorization could be explained in terms of network-level descriptions of brain activity. Our first goal was to focus on graph theoretical approaches to analyze the complex networks that could provide a powerful new way of quantifying individual differences in speech perception. A second goal was to discover which aspects of those functional connectivity networks best explained the variation and diversity in listeners’ perceptual responses during speech sound categorization. We recoded high-density electroencephalograms (EEGs) while listeners rapidly classified speech in a speeded vowel identification task (Bidelman et al., 2013; Bidelman and Walker, 2017). We then applied graph analyses to source-localized EEG responses to derive the underlying functional brain networks related to speech categorization. Using Bayesian non-parametric modeling, we then show that speeded categorical decisions unfold in three RT clusters that distinguish subgroups of listeners based on their behavioral performance (i.e., slow, medium, and fast perceivers). Applying state-of-the-art machine learning and stability selection analyses to neural data we further show that local and global network properties of brain connectomics can decode group differences in behavioral CP performance with 92% accuracy (AUC=0.9). Our findings demonstrate that slow RT decisions related to categorical speech perception involve improper (or excessive) utilization of functional brain networks underlying speech whereas fast and medium perceivers show less utilization.

## METHODS

### Participants

Thirty-five adults (12 male, 23 females) were recruited from the University of Memphis student body and Greater Memphis Area to participate in the experiment. All but one participant was between the age of 18 and 35 years (M = 24.5, SD = 6.9 years). All exhibited normal hearing sensitivity confirmed via audiometric screening (i.e., < 20 dB HL, octave frequencies 250 - 8000 Hz), were strongly right-handed (77.1± 36.4 laterality index (Oldfield, 1971)), and had obtained a collegiate level of education (17.2 ± 2.9 years). None had any history of neuropsychiatric illness. On average, participants had 5.1± 7.5 years of formal music training. All were paid for their time and gave informed consent in compliance with a protocol approved by the Institutional Review Board at the University of Memphis. Figure 1 (A, B) shows the distribution of demographic measures (gender and age) of participants.

**Figure 1:**
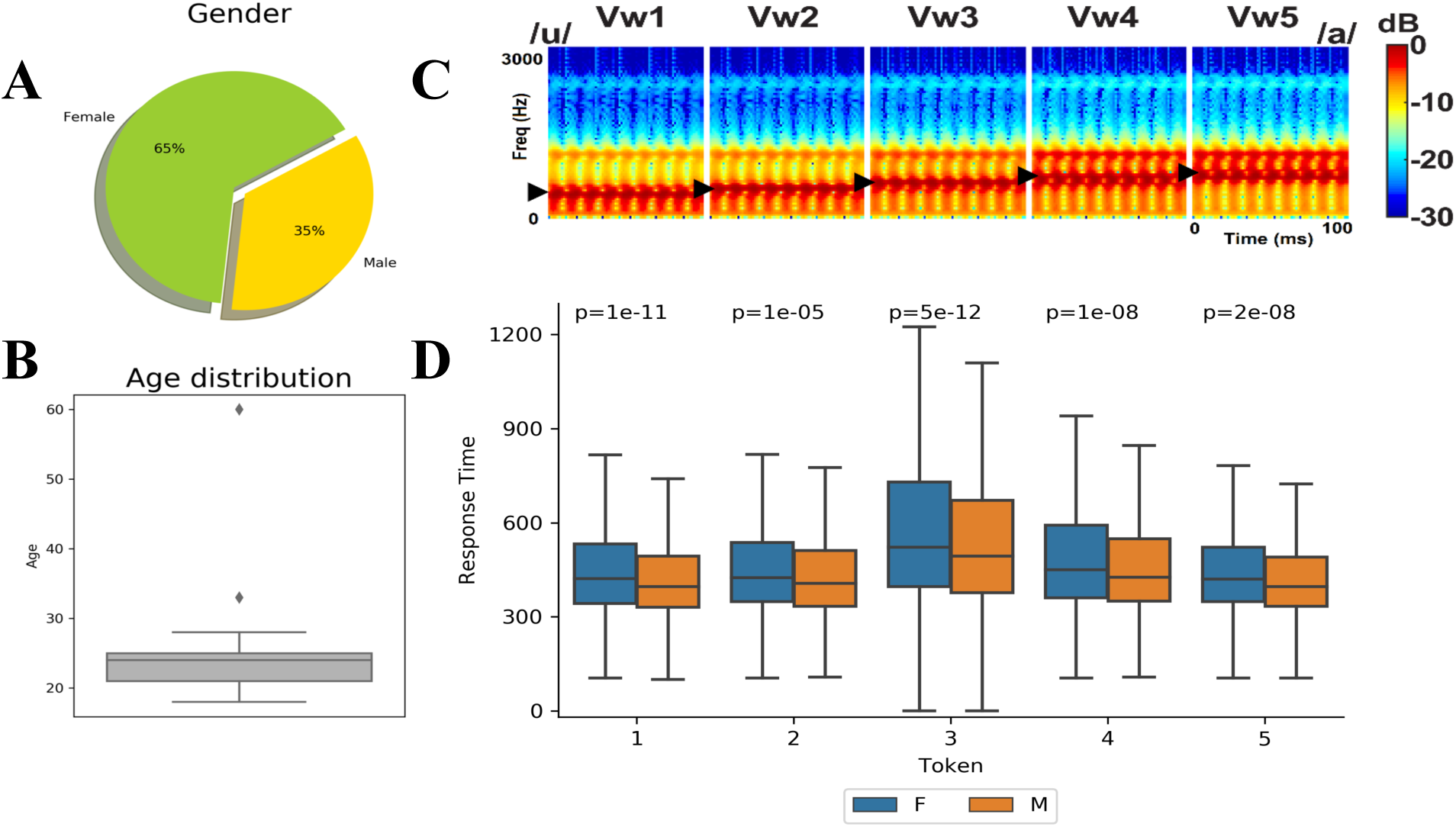
(A, B) Demographic gender and age distributions. (C) Acoustic spectrograms of the speech stimuli: The stimulus continuum was created by parametrically changing vowel first formant frequency over five equal steps from 430 to 730 Hz (▸), resulting in a perceptual-phonetic continuum from /u/ to /a/. (D) Token wise response times for auditory classification. Listeners are slower to label sounds near the categorical boundary (i.e., Token 3). Females (F) have significantly slower response times than males (M).

### Speech stimulus continuum and behavioral task

We used a synthetic five-step vowel continuum previously used to investigate the neural correlates of CP (Oldfield, 1971) (Figure 1C). Each token of the continuum was separated by equidistant steps acoustically based on first formant frequency (F1) yet was perceived categorically from /u/ to /a/. Tokens were 100 ms, including 10 ms of rise/fall time to reduce spectral splatter in the stimuli. Each contained an identical voice fundamental (F0), second (F2), and third formant (F3) frequencies (F0: 150, F2: 1090, and F3: 2350 Hz). The F1 was parameterized over five equal steps between 430 and 730 Hz such that the resultant stimulus set spanned a perceptual phonetic continuum from /u/ to /a/ (Bidelman et al., 2013). Speech stimuli were delivered binaurally at 83 dB SPL through shielded insert earphones (ER-2; Etymotic Research) coupled to a TDT RP2 processor (Tucker Davis Technologies).

During EEG recording, listeners heard 150-200 trials of each individual speech token. On each trial, they were asked to label the sound with a binary response (“u” or “a”) as quickly and accurately as possible (speeded classification task). Reaction times (RTs) were logged, calculated as the timing difference between stimulus onset and listeners’ behavioral response. Following their keypress, the inter-stimulus interval (ISI) was jittered randomly between 800 and 1000 ms (20 ms steps, uniform distribution) and the next trial was commenced.

### Behavioral data analysis

We adopted classical Gaussian mixture modelling (GMM) with expectation-maximization (EM) to identify an optimal number of clusters (i.e., subgroups of listeners) from the distribution of their RT speeds (see Figure 1D). GMMs are probabilistic models that assume the data are generated from a mixture of a finite number of Gaussian distributions (components) with unknown parameters. Mixture models generalize k-means clustering to incorporate information about the covariance structure of the data as well as the centers of the latent Gaussians. Unlike Bayesian procedures, such inferences are prior-free. However, finding an optimal number of components is challenging. The Bayesian Information Criterion (BIC) can be used to select the number of components in a GMM, if data is generated from an i.i.d. mixture of Gaussian distributions. In this study, we used brute-force and BIC based approaches as an alternative solution to the Variational Bayesian Gaussian mixture model. In this exhaustive parameter search, the hyper parameters were (1) Number of components (clusters), (ranges from 1 to 14); (2) Type of covariance parameters (‘full’: each component has its own general covariance matrix; ‘tied’: all components share the same general covariance matrix; ‘diag’: each component has its own diagonal covariance matrix; or ‘spherical’: each component has its own single variance).

This identified an optimal combination four components with unique covariance matrix. Figure 2A shows the BIC scores while tuning parameters. The ‘*’ indicates the optimal combination of components. The probability of each component (see Figure 2B) shows that most trials fall into components 1-3 ranging from 17% - 47% of the total trials in the speech identification task. Component 4 has the fewest number of trials (1.6%). Based on the interpretation of RTs, we categorized these components as Fast RT (Cluster 2, 120 - 476 ms), Medium RT (Cluster 3, 478 - 722 ms), Slow RT (Cluster 1, 724 -1430 ms), and Outliers (Cluster 4, 1432 - 2500 ms). The outliers (Cluster 4) were discarded for further analysis given the low trial counts loading into this cluster. The boxplot in Figure 2C shows token wise response times. Each speech token can be broken down into a combination of the three RT clusters, meaning that speech categorization speeds could be objectively clustered into fast, medium, slow (and outliers) responses via the GMM. These cluster divisions were then used in subsequent EEG analyses to determine if functional brain connectomics differentiated these subgroups of CP performers.

**Figure 2:**
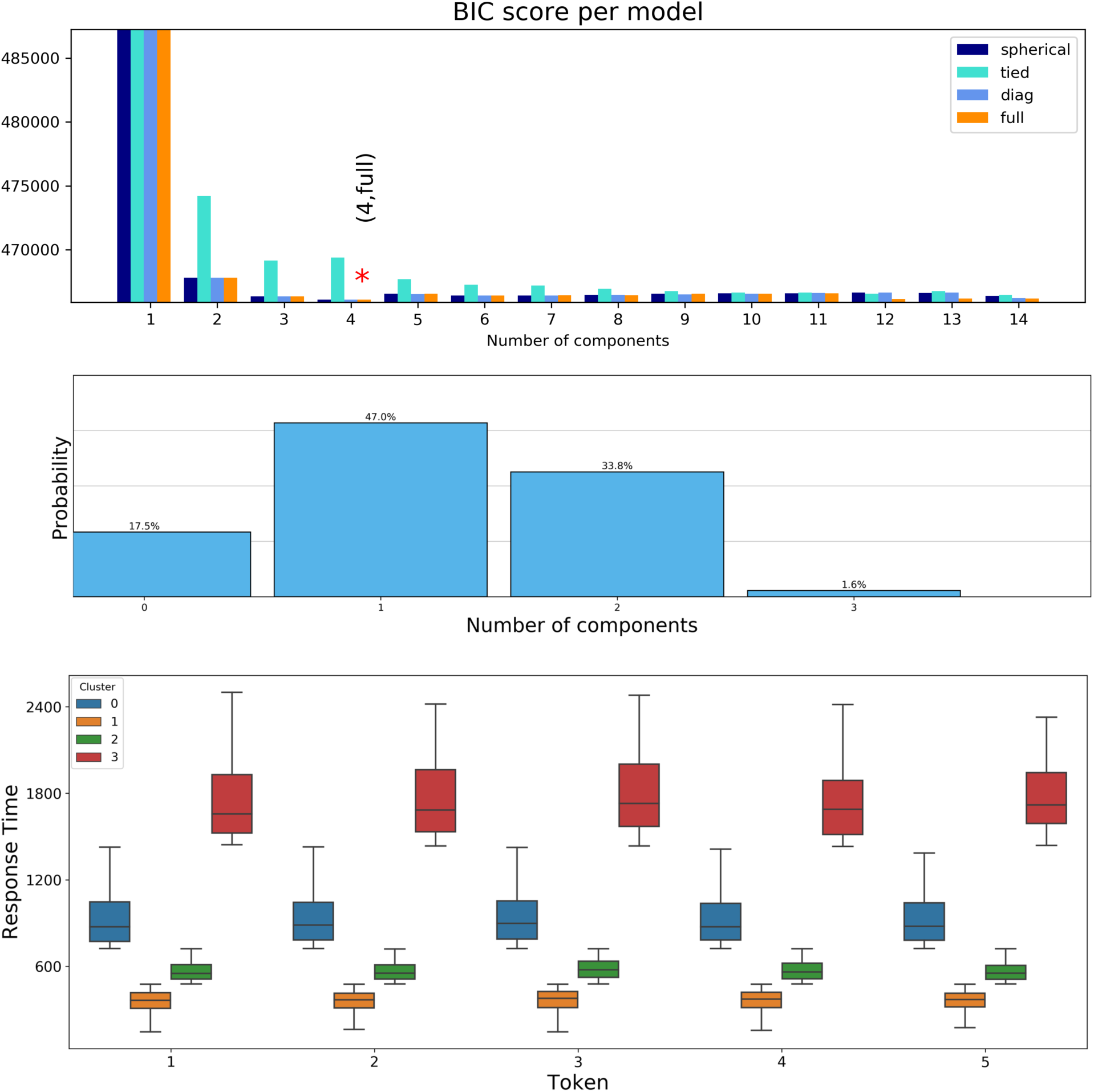
Clustering RT data using GMM and BIC criteria. Model selection concerns both the covariance type and number of components in the model. Brute-force based empirical analysis shows that n=4 components with unique covariance matrix is optimal. The ‘*’ marked position of (A) shows the optimal combination. (B): Probability of trials loading into each component. (C): Token wise RT broken down by component. Based on behavioral RTs, four clusters are evident that distinguish subgroups of listeners based on their speech identification speeds: Fast (Cluster 1): 120∼476 ms, Medium (Cluster 2): 478∼722 ms, Slow (Cluster 0): 724∼1430 ms, and Outliers (Cluster 3): 1432∼2500 ms.

### EEG recording and preprocessing

#### Recording and preprocessing

EEG recording procedures were identical to our previous neuroimaging studies on CP (e.g., (Bidelman et al., 2013; Bidelman and Alain, 2015; Bidelman and Walker, 2017)). Briefly, neuroelectric activity was recorded from 64 sintered Ag/AgCl electrodes at standard 10-10 locations around the scalp (Oostenveld and Praamstra, 2001). Continuous data were digitized using a sampling rate of 500 Hz (SynAmps RT amplifiers; Compumedics Neuroscan) and an online passband of DC-200 Hz. Electrodes placed on the outer canthi of the eyes and the superior and inferior orbit monitored ocular movements. Contact impedances were maintained < 10 kΩ during data collection. During acquisition, electrodes were referenced to an additional sensor placed ∼ 1 cm posterior to the Cz channel.

Subsequent pre-processing was performed in BESA® Research (v7) (BESA, GmbH). Ocular artifacts (saccades and blinks) were first corrected in the continuous EEG using a principal component analysis (PCA) (Picton et al., 2000). Cleaned EEGs were then filtered (bandpass: 1-100 Hz; notch filter: 60 Hz), epoched (−200-800 ms) into single trials, baseline corrected to the pre-stimulus interval, and re-referenced to the common average of the scalp. This resulted in between 750 and 1000 single trials of EEG data per subject (i.e., 150-200 trials per speech token).

#### Source analysis

Following our previous neuroimaging studies on speech processing(Bidelman and Dexter, 2015; Bidelman and Howell, 2016), we performed a distributed source analysis to more directly assess the neural generators underlying behavioral decisions related to CP. Source reconstruction was implemented in the MATLAB package Brainstorm (Tadel et al., 2011). We used a realistic, boundary element model (BEM) volume conductor (Fuchs et al., 2002, 1998) standardized to the MNI brain (Mazziotta et al., 1995). The BEM head model was created using the OpenMEEG (Gramfort et al., 2010) as implemented in Brainstorm (Tadel et al., 2011). A BEM is less prone to spatial errors than other head models (e.g., concentric spherical conductor) (Fuchs et al., 2002). sLORETA allowed us to estimate the distributed neuronal current density underlying the measured sensor data. The resulting activation maps (akin to fMRI) represent the transcranial current source density underlying the scalp-recorded potentials as seen from the cortical surface. We used the default settings in Brainstorm’s implementation of sLORETA (Tadel et al., 2011). From each single-trial sLORETA map, we extracted the time-course of source activity within 68 regions of interest (ROI) defined by the Desikan-Killany Atlas parcellation (Desikan et al., 2006) as implemented in Brainstorm. Single-trial source waveforms (derived per subject and speech token) were then submitted to functional connectivity analyses. We have recently used a similar approach to successfully decode single-trial EEG and predict individual differences in other cognitive domains (e.g., working memory capacity (Bashivan et al., 2017)), motivating its use here.

### EEG functional connectivity and graph analyses

#### Bootstrapping

Functional connectivity measures are more accurate when calculated using source localized compared to scalp-recorded (sensor-level) EEG (Brunner et al., 2016). Still, to ensure robustness of our connectivity measures, we used bootstrapping to reduce the uncertainty of our connectivity estimates (James et al., 2013). This method involved repeatedly taking small samples with replacement, calculating the statistics, and averaging over the calculated statistics. We applied a mean based bootstrap approach on 35106 trials. For each RT class, 100 random trials from each individual participant were chosen as a bootstrap sample (with replacement). We calculated the mean source amplitude in each of the 68 ROIs for each bootstrap sample. This process was then iterated 30 times to derive the final estimate of the mean source signal in each ROI. Overall, 3150 trials were generated (1050 trials of each RT class) in this process for further analysis.

#### Functional connectivity

A graph network is defined by a collection of nodes (vertices) and links (edges) between pairs of nodes. Nodes in large-scale brain networks usually represent brain regions (ROIs), while links represent anatomical, functional, or effective connections (Friston et al., 1994). Anatomical connections typically correspond to white matter tracts between pairs of brain regions. However, functional connections correspond to the strength of temporal correlations between pairs of anatomically connected/unconnected regions. Depending on the measure, functional connectivity may reflect linear or nonlinear interactions, as well as interactions at different time scales (Zhou et al., 2009). To quantify functional connectivity, we measured pair-wise Pearson product-moment correlation coefficients among the 68 brain regions (ROIs). This resulted in connectivity matrix describing the weighted strength (undirected network) between all pairwise nodes (^68^C_2_ = 2278 edges) for each trial. Diagonal and upper diagonal elements of the connectivity matrices were discarded to avoid spurious self and repeated connectivity. Matrices were then concatenated to a vector to describe the connectivity across all brain nodes and trials (e.g., 3150*2278) for each participant.

Seven global network connectivity features were estimated from each network graph using the BCT toolbox (Rubinov and Sporns, 2010): (i) Characteristics path, (ii) Global efficiency, (iii) Average clustering coefficient, (iv) Transitivity, (vi) Small-worldness, (vi) Assortativity coefficient, and (vii) Maximized modularity (see Appendix for mathematical definitions and interpretation of these network features).

### Machine learning: identifying behaviorally-relevant aspects of functional connectivity

The t-distributed stochastic neighbor embedding (t-SNE) (Maaten and Hinton, 2008) is a widely used unsupervised learning algorithm to visualize high-dimensional data. t-SNE converts similarities between higher dimensional data points to joint probabilities, providing a faithful representation of those data points in a lower-dimensional human interpretable 2D or 3D plane. Such a projection brings insight on whether the data is separable, the data lies in multiple different clusters, or inspecting the nature of those clusters. We adopted LDA on our three-class connectivity dataset (i.e., fast, medium, slow perceivers identified from the behavioral data) and considered 50 dimensions for t-SNE. The hyper parameters of t-SNE were tuned with a grid search approach. Figure 3 shows the t-SNE embedded scatter and kernel density estimation (KDE) plot of our data distribution. KDE plot is a non-parametric way to represent the probability density function and is used here to visualize the trend of the data distribution for each different class (data points for fast, medium, and slow RTs). The t-SNE visualization shows three nearly distinct clusters of functional connectivity for the different RT groups in speech categorization. Unrelated or noisy edges may exist in the higher dimensional functional connectivity matrices. This necessitates the use of feature selection methods to choose functional connectivity metrics that are relevant and can be modeled robustly over a range of model parameters.

**Figure 3:**
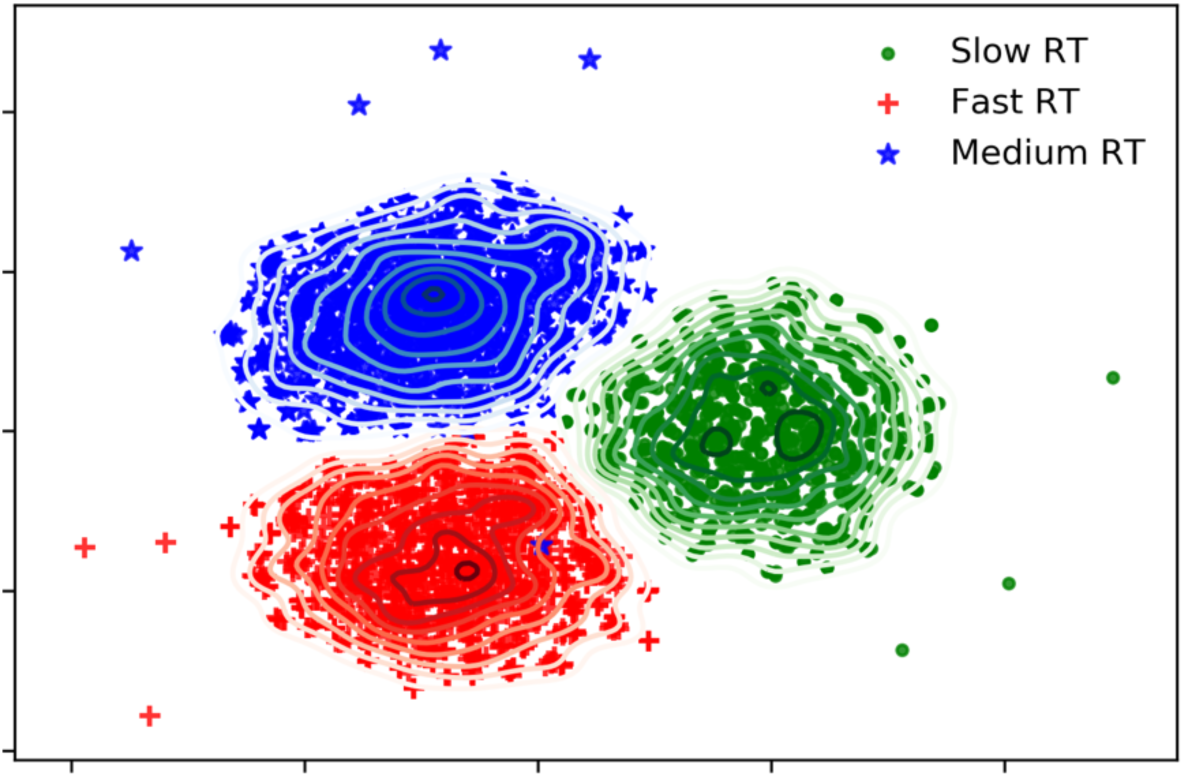
The t-SNE embedded higher dimensional functional connectivity data are represented by a 2-dimensional scatter and kernel density estimation (KDE) plot. The green lines with ‘.’, blue lines with ‘*’, and red lines with ‘+’sign represents data points for slow, medium, and fast RT participants, respectively.

### Feature selection (stabilty selection)

Feature selection attempts to identify the most salient subset of variables from a larger set of features mixed with irrelevant variables. This problem is especially challenging when the number of available data samples are smaller compared to the number of possible features. Conventional filter methods identify a consistent set of variables outside of the predictive model based on some filtering criteria, e.g., the variables are individually evaluated to check the probable relationship between classes. The sets of variables in this technique are selected based on a threshold of importance. Commonly filter-based methods include correlation, F-test, chi-square test, ANOVA analysis. The highly-correlated or redundant features may be selected, and significant interactions and relationships between variables may not be able to be quantified. However, one of the downsides of the multivariate approaches (e.g., PCA, LDA, Lasso, Elasticnet SVM ranking, Wrapper based methods, GA Wrapper, Forward Backward based methods) is that outcomes often depend on model parameters (e.g., regularization factor). Compared to conventional filter and multivariate approaches, stability selection produces more reliable estimations and yields a stable set of features because of its internal randomization implemented as bootstrap based subsampling. It was reported that even if the necessary conditions needed for consistency of the original Lasso (L1 norm penalized linear models) method are violated, stability selection will be consistent in variable selection (Meinshausen and Bühlmann, 2010). The main advantages of this algorithm are (1) it works efficiently with the high-dimensional data, (2) stability selection provides finite sample control with error rates of false discoveries and is a transparent method to choose a amount of regularization for structure estimation; and (3) it is extremely general and has a very wide range of applicability.

An attractive feature of Lasso (L1 regularization on least squares) is its computational feasibility for the high-dimensional data with many more variables than samples since the optimization problem of lasso estimator is convex. Furthermore, the Lasso can select variables by shrinking certain estimated coefficients exactly to 0. Hence Lasso was used for stability selection. Applying Randomized Lasso many times and looking for variables that are chosen is a very powerful procedure tool to select consistent or stable features (Al-Fahad et al., 2017; Meinshausen and Bühlmann, 2006; Shah and Samworth, 2013; Tibshirani, 1996). Despite its simplicity, it is consistent for variable selection even though the ‘neighborhood stability’ condition is violated. More about stability section, interpretation and mathematical definition are explained in the appendix.

We used Randomized Logistic Regression for stability selection with randomized lasso. It works by subsampling the training data and fitting an L1-penalized Logistic Regression model where the penalty of a random subset of coefficients has been scaled. We considered sample fraction = 0.75, number of resampling =1000 with tolerance=0.001. This algorithm assigns feature scores between 0 and 1 based on frequency of selection over 1000 iterations. We need to specify the score to find out the best representative set of stable features. Hence, threshold selection is a design parameter. We varied different selection thresholds (i.e., the number of selected features) and observed the effect on model performance. Modeling involved four steps:

1. Randomly shuffle and split the dataset in to training and test set (80% and 20%).

2. Consider Support Vector Machine with “RBF” kernel as a base estimator.

3. Tune hyper parameter (i.e. C and Gamma) on training data using grid search approach and10-fold cross validation.

4. Selected best models are evaluated on unseen test data. Accuracy (ACC) and Area Under Curve (AUC) were considered for performance measure.

Figure 4 shows the effect of different selection thresholds on modeling. The histogram illustrates the distribution of the feature score. The first line of the x axis shows the bin ranges of scores (0 to 1. The second and third lines show the amount and percent of features that had nearly the same score for a specific bin. We found 73% of the features had scores of 0-0.1, meaning the majority of connectivity measures were not selected even once (i.e., coefficient was zero) among 1000 model iterations. That is, 73% of functional connectivity metrics explored in our search space were not related to speeded speech categorization (i.e., behavioral RTs).

**Figure 4:**
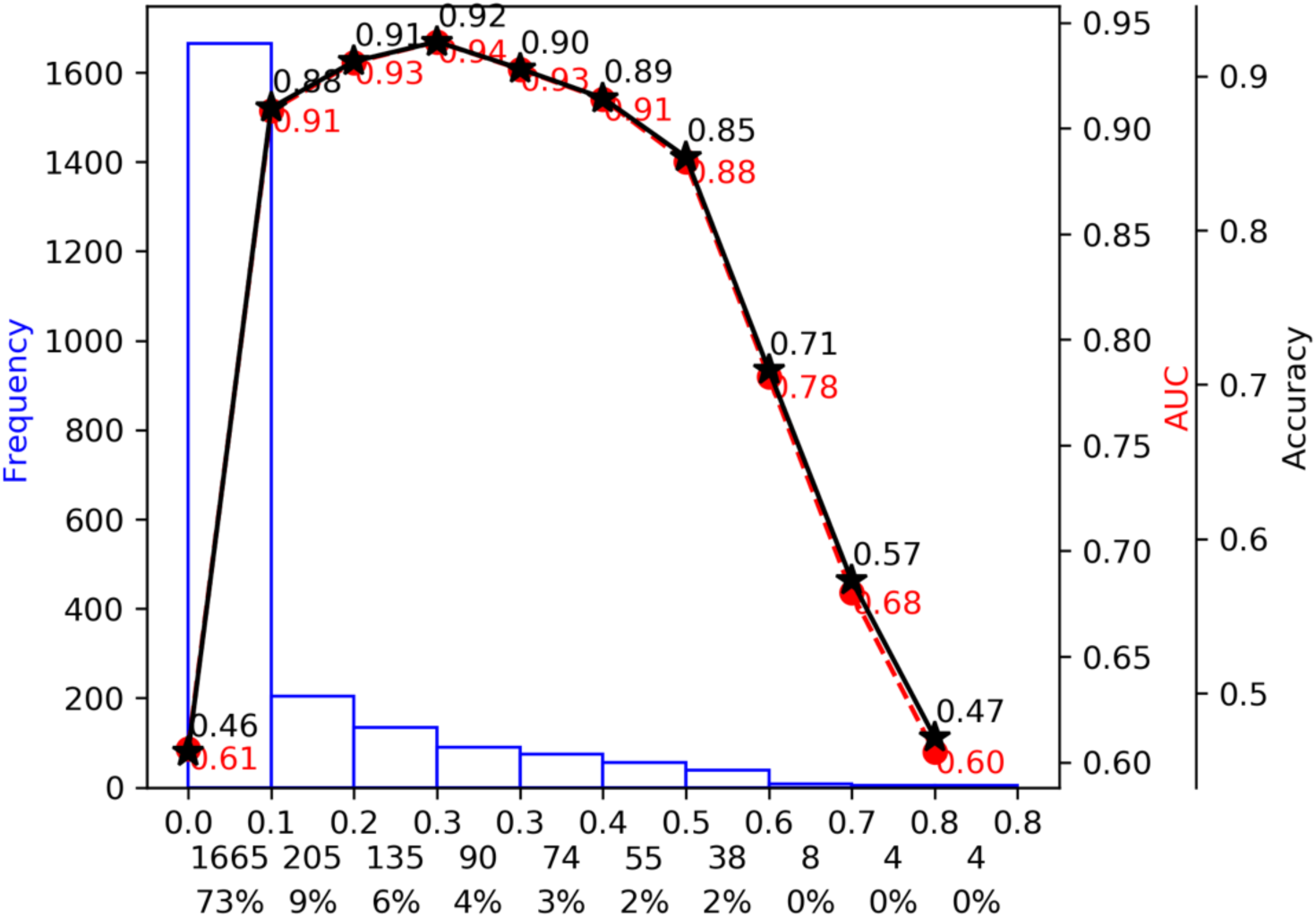
Effect of selection threshold on model performance prediction. The three x-labels represent (top) the range of each bin of features score (range: 0∼1), (middle) the number of features falling in each bin, and (bottom) the corresponding percentage.

For a specific selection threshold of 0.26, the algorithm selected 227 edge features that collectively achieved 92% accuracy (best model performance) with AUC=0.9. The bell shaped solid black and red dotted lines of Figure 4 shows the Accuracy and AUC curves for different selection thresholds. Note that selection thresholds higher than the optimal value (0.26) allowed the model to consider more noise variables, degrading model performance significantly. On the other hand, selection thresholds higher than the optimal value discard behaviorally relevant features and reduce model performance. Table 3 details the effect of selection threshold on model performance. Here, the number of unique edges represents correlation-based connectivity between two brain nodes (features) and the number of unique nodes represents brain regions associated with those selected edges. A schematic diagram of the method pipeline is shown in Figure 5.

**Figure 5:**
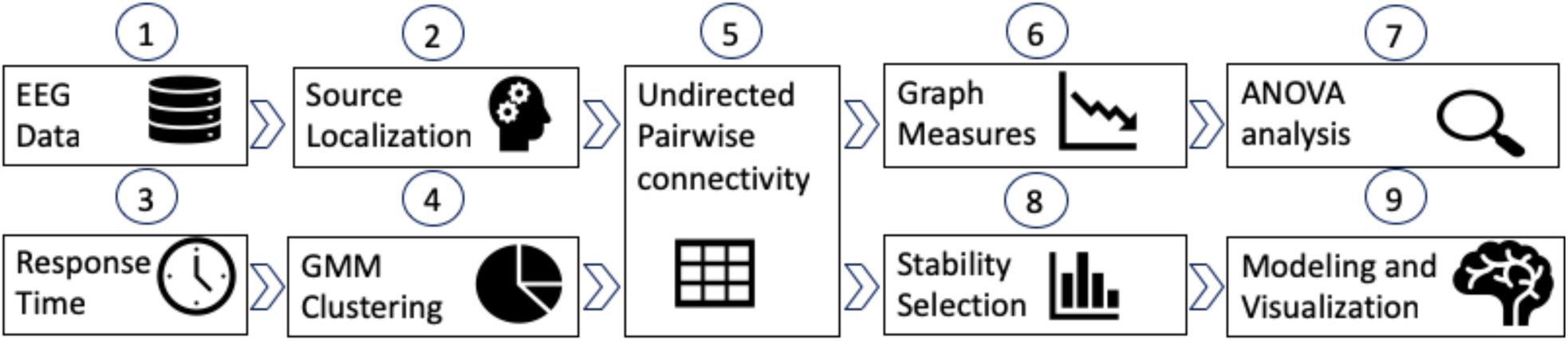
Schematic diagram of the processing pipeline. The 64-ch EEG data is first preprocessed, and then source localization is adapted to convert skull surface data to cortical surface time series data (68 ROIs defined by the Desikan-Killany Atlas parcellation). Pairwise correlations were calculated to derive the connectivity matrix for each trial of the speech CP task. Behavioral response times (RTs) were clustered with Bayesian non-parametric (GMM) clustering. These clusters were labeled as Fast, Medium and Slow RT. ANOVA analysis of Graph measures w adopted to test significance among RT groups. Stability selection and machine learning approaches were then used to find significant properties of the brain’s functional connectivity related to behavioral speeds (RTs) in speech CP.

## RESULTS

Figure 1D shows behavioral results in the speech categorization task. Generally speaking, listeners were slower to label sounds near the categorical boundary (token 3), consistent with the higher ambiguity of the mid-continuum stimuli (Bidelman et al., 2013; Liebenthal et al., 2010; Pisoni and Tash, 1974; Reetzke et al., 2018). On average, females also showed slower RTs than males across the continuum (Welch’s t-test; p<0.0001). Bayesian nonparametric clustering revealed four distinguishable clusters in the speed (RTs) of listeners’ CP (Fast: 120∼476 ms, Medium: 478∼722 ms, Slow: 724∼1430 ms, and Outliers: 1432∼2500 ms) (Figure 2C). These clusters were even present at the individual token level.

Having established that listeners could be distinguished based on their speed in speech categorization, our next goal was to determine whether network properties of the brain accounted for these behavioral differences. We applied graph theory techniques to construct and analyze the functional brain connectome underlying CP. We considered both individual trial- as well as group-based analyses. For group-based analysis, data were averaged across subjects within each RT cluster. Group means were computed by concatenating group-wise trials and calculating their mean. We then calculated seven global network connectivity features using the BCT toolbox (Rubinov and Sporns, 2010) (see Methods).

We used non-parametric ANOVAs (Kruskal-Wallis H-test) to determine if individual trial-based global graph measures varied across RTs (Table 1) This non-parametric test was used given the unequal sample size per group (Lowry, 2014). These analyses revealed that Assortativity and Global Efficiency were modulated depending on behavior speed. Table 2 shows a comparison of the graph measures across three RT groups. Global efficiency measures were relatively small, and assortativity had a negative tendency. All other network features were not discriminatory among the RT groups. Therefore, modeling with those features (using SVM with ‘RBF’ kernel described in method section) showed expectedly poor accuracy (38%).

**Table 1:** Significant (bold) global network measures (Kruskal-Wallis H-test tests) (trial-level)

| Measures | p-value |
| --- | --- |
| Characteristics Path | 0.1359 |
| Average Clustering Coefficient | 0.8286 |
| Small Worldness | 0.0815 |
| **Assortativity** | **0.0052** |
| **Global Efficiency** | **0.0290** |
| Transitivity | 0.8424 |
| Maximized Modularity | 0.6617 |

**Table 2:**
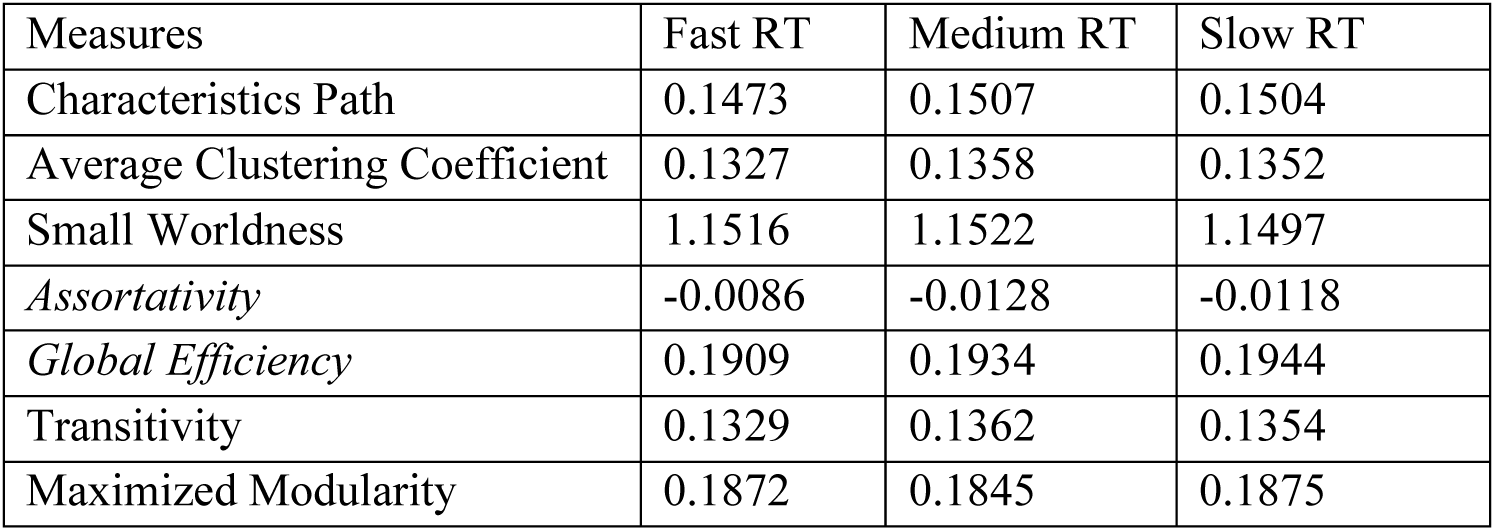
Group comparison of graph measures of functional connectivity between RT groups.

**Table 3:** Effect of selection threshold of stability selection (Threshold) on model performance. Pairwise correlation between two brain regions (functional connectivity edge) were consider as features. Number of unique nodes are brain regions associated with selected features. ACC, accuracy; AUC, area under curve.

| Threshold | ACC | AUC | Number of Unique Edge (features) | Number of Unique Node |
| --- | --- | --- | --- | --- |
| 0 | 46% | 0.6 | 2278 | 68 |
| 0.08 | 88% | 0.9 | 613 | 68 |
| 0.17 | 91% | 0.9 | 408 | 68 |
| 0.26 | 92% | 0.9 | 273 | 68 |
| 0.34 | 90% | 0.9 | 183 | 68 |
| 0.42 | 89% | 0.9 | 109 | 64 |
| 0.51 | 85% | 0.9 | 54 | 53 |
| 0.59 | 71% | 0.8 | 16 | 24 |
| 0.68 | 57% | 0.7 | 8 | 13 |
| 0.76 | 47% | 0.6 | 4 | 8 |

Besides analyzing global network properties, we next aimed to identity the most significant properties of functional brain connectivity that were related to behavioral RTs. Functional connectivity for each trial is a high dimensional sparse matrix. Some studies have suggested that properties of functional brain networks are most consistent with the actual brain anatomy when network density is 8–16% (Li et al., 2016; Salvador et al., 2005; Wang et al., 2010). To determine the most behaviorally-relevant arrangement of sparse connectivity, we used stability selection with Randomized Lasso to detect and rank the most important, consistent, and relevant functional connectivity measures that were invariant (stable) over a range of model parameters. Stability selection discarded 88% (total 273) of network edges that were not related to behavioral RTs, but still achieved 92% classification accuracy with AUC=0.9. From Table 3, It was observed that only 7% error tolerance from the optimal value (accuracy from 92% to 85%) allow 80% less edge and 22% less associated nodes. Hence, the selection threshold 0.51 with reasonable performance (ACC=85%, AUC=0.9) were chosen for network visualization as performance declined precipitously above this threshold (Figure 4).

Figure 6 shows a visualization of the 54 nodes among 53 ROIs identified via stability selection using BrainNet (Xia et al., 2013). The resulting network revealed a highly dense connectome reflective of listeners’ behavioral RTs in speech categorization. Connectivity was particularly strong between the occipital, parietal, and bilateral frontal lobes. As an additional means of data reduction, we further threshold (=0.68) the stability-selected connectome. This resulted in eight highly ranked connectivity edges among 13 nodes across the brain (Figure 7). Even with this sparse network of only 8 edges, model classification was still 57%, meaning this small set of features accuracy predicted RTs. We then ranked the contribution of these stable nodes that described behavior Table 4. We found that three edges (rank: 3, 4, and 6) were in left hemisphere, two edges were in right hemisphere (rank: 2, and 5) and three edges were inter-hemispheric (rank: 1, 7 and 8). Notably, these edges included connections between motor (paracentral), visual (lateral occipital/ lingual), linguistic (pars triangularis), auditory (superior temporal gyrus), and parietal areas both within and between hemispheres.

**Table 4:** Eight most important edges that govern speeded speech classification. Collectively, these edges achieve a model accuracy of 57% in segregating listeners’ speeded decisions (RTs) in the perceptual task. Here, a score of 0.85 means that out of 1000 iterations, the edge was selected by stability selection 850 times.

| Edge | Score | Rank |
| --- | --- | --- |
| Paracentral R-Middletemporal L | 0.85 | 1 |
| Lingual R-Caudalmiddlefrontal R | 0.845 | 2 |
| Parstriangularis L-Inferiorparietal L | 0.785 | 3 |
| Superiorparietal L-Rostralmiddlefrontal L | 0.785 | 4 |
| Precuneus R-Parahippocampal R | 0.725 | 5 |
| Parstriangularis L-Lateraloccipital L | 0.705 | 6 |
| Precuneus R-Lingual L | 0.705 | 7 |
| Superiortemporal R-Inferiorparietal L | 0.695 | 8 |

**Figure 6:**
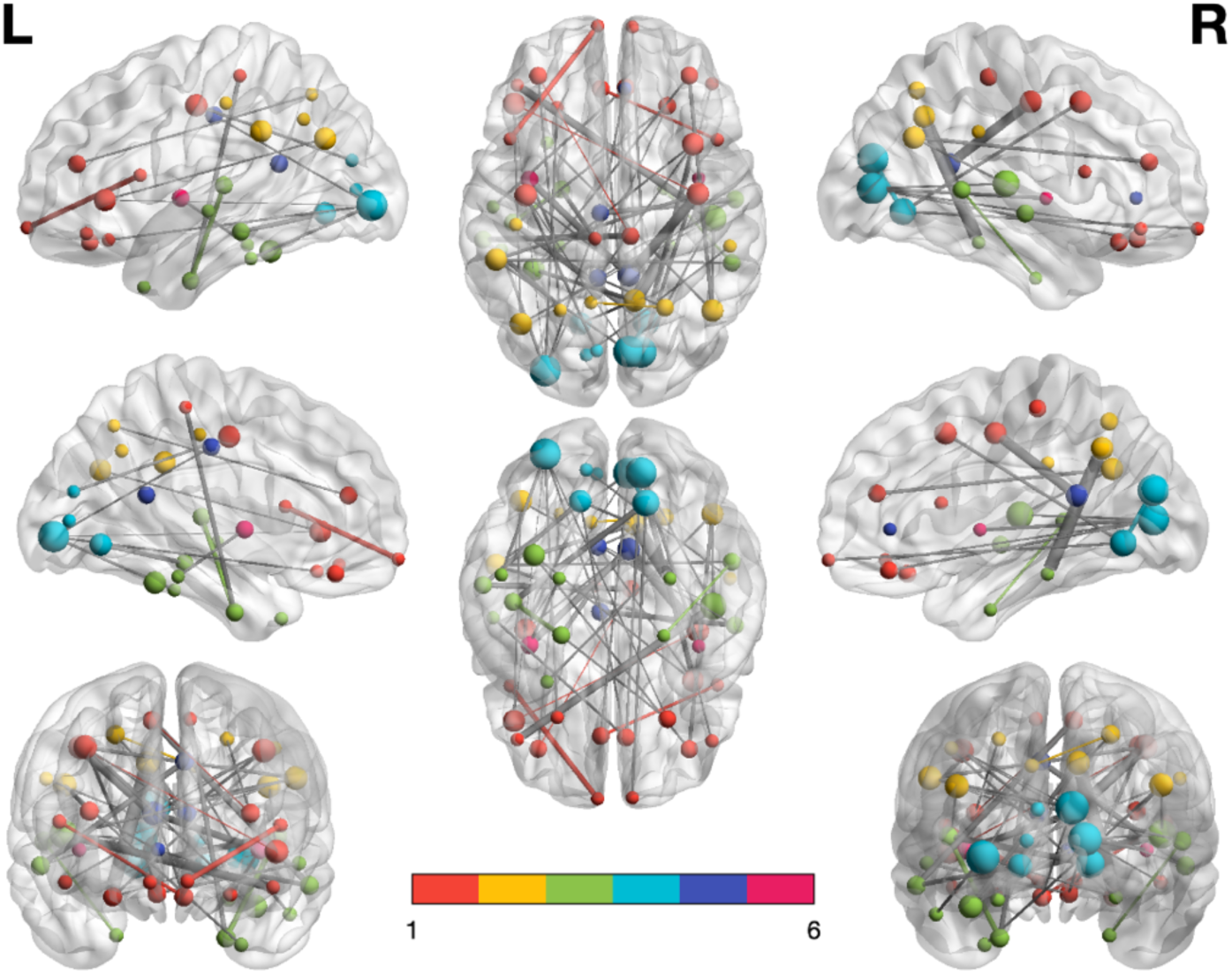
BrainNet visualization (top to bottom: lateral, medial, and dorsal view) of the brain network (54 edges) identified via stability selection. Color map 1-6 indicates, 1: Frontal (22 ROI), 2: Parietal (10 ROI), 3: Temporal (18 ROI), 4: Occipital (8 ROI), 5: Cingulate (8 ROI), 6: Insula (2 ROI) regions. Node size varies with its degree of connectivity. Connectivity among the same lobe are colored with similar node color. Edge widths represent the weight of absolute correlation (connectivity strength).

**Figure 7:**
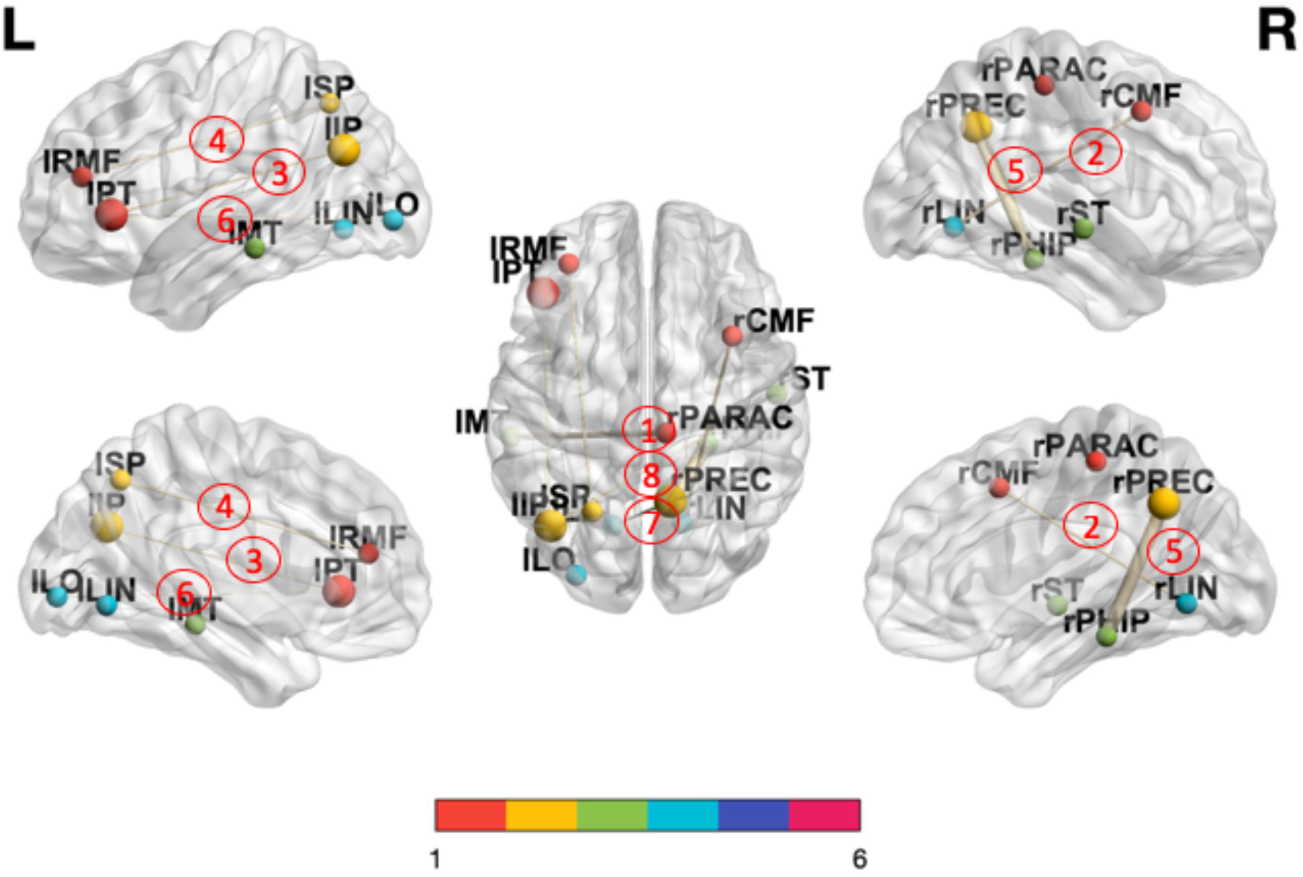
A sparse brain network (8 edges) predicts listeners’ speed (RTs) of speech categorization (57% model accuracy). Red numbers are the ranked importance of the edges describing behavior. Otherwise as in Figure 6.

**Figure 8:**
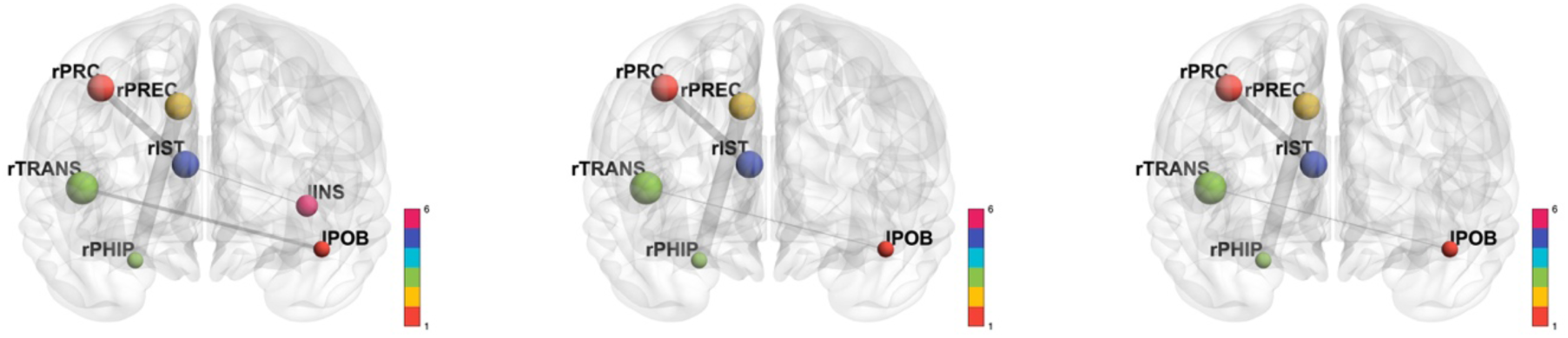
Brain network underlying Slow RT listeners (left), Medium RT listeners (middle), and Fast RT listeners. Shown here are the most highly correlated (absolute correlation ≥0.5) network edges. Otherwise as in Figs. 6-7. INS, insula; IST, isthmus of cingulate; TRANS, transverse temporal gyrus (auditory cortex); POB, pars orbitalis; PRC, precentral gyrus (motor cortex); PHIP, parahippocampal gyrus; PREC, precunus; l/r, left/right hemisphere.

## DISCUSSION

The present study evaluated whether individual differences in a core operation of speech and language function (i.e., categorization) could be explained in terms of network-level descriptions of brain activity. By applying machine learning classification techniques to functional connectivity data derived from EEG, our data show that the speed of listeners’ ability to categorize and properly label speech sounds is directly related to dynamic variations in their brain connectomics.

It has been suggested that important cognitive functions are supported by distributed neural networks with highly segregated and integrated “small-world” organizations or clusters (Bassett and Bullmore, 2006; Honey et al., 2007; Newman, 2003; Tononi et al., 1994). However, in relation to distinguishing listeners’ perceptual speed for categorized speech, we did not find differences in network properties of Characteristics Path, Average Clustering Coefficient, Small Worldness, Transitivity and Maximized Modularity clearly indicates (Table 1 and Table 2). Instead, global network assortativity and efficiency distinguished fast, medium, and slow RT individuals. In network science, assortativity refers to the tendency of “like to connect with like.” That is, at the macroscopic level, high degree nodes attach to other high degree nodes and similarly, low to low (Stam et al., 2014). In our study, functional brain networks were defined via task-based co-activations. Hence, they were expected to exhibit some assortativity as co-activation means that regions of the network were engaged by the same task. Previous studies have shown that the property of assortative tendency changes with task demands (Betzel et al., 2018). The resting state brain functional network is largely assortative. Higher order association areas exhibit non-assortative organization tendency and form periphery-core topologies. However, assortative structures break down during tasks and is supplanted by periphery, core, and disassortative communities.

In addition, we found that the functional CP network underlying speeded decisions increased in negative assortativity (i.e., became disassortative) for slower RTs (Table 2). This indicates that brain nodes were more likely to connect with nodes having different degree during slower RTs, implying that important hubs of the CP network communicated with insignificant hubs during states of slower decisions. Based on the interpretation of these graph metrics (see Appendix), we infer that slower, more taxing categorical speech decisions cause excessive use of neural resources and more aberrant information flow within the CP circuitry. Supporting this interpterion, we found that Network utilization (Global efficiency) also differentiated RT groups. Higher Global efficiency indicates that the routing of information among nodes with different degree was significantly higher for slow RT trials. In short, we find that slower perceivers tended to utilize functional brain networks excessively (or inappropriately) whereas fast perceivers (with lower global efficiency) utilized the same neural pathways but with more restricted organization. Presumably, these dynamic changes in brain connectivity account for the variations in RTs we find during speech categorization at the behavioral level (Figure 1D).

Our data show that global graph measures fail to fully explain the behavioral relevance of important connectivity edges. We observed the functional connectivity matrix underlying speech CP is highly sparse and dynamic. Indeed, only ∼12% of all possible edges in the Desikan-Killany Atlas were needed to explain variation in behavioral RTs. In this vein, we adopted stability selection to find edges that were most consistent in distinguishing different network states related to perception. By performing this two stage of randomization iteratively (e.g., 1000 bootstraps), stability selection with randomized lasso assigned high scores to features that were repeatedly selected across randomizations, yielding the most meaningful connections within the CP connectome that describe behavior.

Collectively, our results showed that neural classifiers (SVM) coupled with stability selection can correctly classify behavioral RTs related to CP from functional connectivity alone with over 90% accuracy (AUC=0.9). The resulting edges composing the RT-related networks were distributed in both hemispheres and both intra- and inter-hemispheric edges were evident. More interestingly, we found that only 8 edges among 13 ROIs were needed to distinguish RTs well above chance (Figure 7). ROIs composing this sparse but behaviorally-relevant network included (1) Caudalmiddlefrontal R, (2) Inferiorparietal L, (3) Lateraloccipital L, (4) Lingual L, Lingual R, (6) Middletemporal L, (7) Paracentral R, (8) Parahippocampal R, (9) Parstriangularis L, (10) Precuneus R, (11) Rostralmiddlefrontal L, (12) Superiorparietal L, (13) and Superiortemporal R. Previous neuroimaging studies have demonstrated a distributed fronto-temporo-parietal neural network supporting auditory categorization (e.g., (Alho et al., 2016; Bidelman and Lee, 2015; Binder et al., 2004; Chang et al., 2010; Feng et al., 2018; Golestani et al., 2002; Golestani and Zatorre, 2004; Lee et al., 2012; Liebenthal et al., 2010; Luthra et al., 2019; Myers et al., 2009)). Our data corroborate these previous studies by confirming engagement of similar temporal (STG), parietal, motor, and prefrontal regions in CP using an entirely data driven approach (machine learning with stability selection).

Notably, we found functional connectivity between right paracentral and left middletemporal gyrus (MTG) was the most important connection describing the speed of behavioral CP (Table 4). MTG has been associated with accessing word meaning while reading (Acheson and Hagoort, 2013) and has been described as an early lexical interface that is heavily involved in sound-to-meaning inference (Hickok and Poeppel, 2007, 2004). Some studies indicate that lesions of the posterior region of the middle temporal gyrus, in the left cerebral hemisphere, may result in certain forms of alexia and agraphia (Sakurai et al., 2008), indicating its role in the language production network (Blank et al., 2002). The strong link between MTG and paracentral gyrus implies a direct pathway between the neural substrates that map sounds to meaning and sensorimotor regions that execute the motor command and therefore govern response speeds (indexed by RTs). The leftward laterality of the MTG node is consistent with the left lateralized nature of language processing in the brain. Still, why left MTG so strongly interfaces with *right* motor areas in our data is unclear, especially given the right-handedness of our participants and expected left (contralateral) motor involvement. Differences in brain connectivity have been observed between sexes (Ingalhalikar et al., 2014) and females may have a more diffuse, bilateral neural system for language processing than males (Shaywitz et al., 1995). Speculatively, the strong communication between left linguistic (MTG) and right motor brain areas we find may reflect the higher preponderance of females in our sample.

Relatedly, stability selection identified the second ranked edge between lingual and caudal-middlefrontal gyrus. While the functional role of lingual (occipital) gyrus in speech processing is not apparent prima facie, this region is involved in visual word processing, especially letters (Mechelli et al., 2000). It has also implicated in stimulus naming (Bookheimer et al., 1995; Howard et al., 1992), an operation at the core of our speech categorization (i.e., sound labelling) task. We also found a third ranked edge predictive of behavioral CP between parstriangularis and inferior parietal cortex. Previous functional neuroimaging and connectivity studies have shown strong engagement of frontal-parietal networks during CP (Feng et al., 2018; Liebenthal et al., 2010; Luthra et al., 2019). Our results corroborate these findings by similarly implicating a strong interface between linguistic (IFG) and parietal (IPL) brain regions in modulating the speed of listeners’ categorical decisions. Indeed, decision loads IFG during effortful speech listening (Binder et al., 2004; Bouton et al., 2018; Du et al., 2014) and the IFG-IPL pathway is upregulated when speech material is perceptually confusable (Feng et al., 2018). Therefore, the network organization of brain connectivity observed for slow RTs and importance of IFG-IPL in describing behavior may reflect a similar state of perceptual confusion during rapid categorical speech labeling.

One limitation of our study was that our sample contained more females than males (2:1 ratio). This is relevant since RTs were significantly different among genders (Figure 1D). Thus, a natural question that emerges from our data is the degree to which our machine learning techniques segregated data based on gender rather than different RTs (i.e., fast vs. slow perceivers), per se. Still, this is probably not the case. Conventional filter-based group analysis can bias classification and feature selection results whereas with our Lasso-based bootstrapped analysis, this becomes less likely (Bach, 2008). Moreover, stability selection with randomized lasso is a similar but more robust approach that produces consistent variable selection with minimal bias. Hence, the impact of our unbalanced sample size on feature selection is probably negligible.

Taken together, our novel approach to neuroimaging data demonstrates the derivation of small, yet highly meaningful patterns of brain connectivity that dictate speech behaviors using solely EEG. More broadly, the functional connectivity and machine learning techniques used here could be deployed in future studies to identify the most meaningful changes in spatiotemporal brain activity that are modulated by development, normal learning, or those which decline in neuropathological states.

## CONCLUSION

We developed an efficient computational framework to investigate whether individual differences in speeded speech categorization could be explained in terms of network-level descriptions of functional brain connectivity. Our EEG data-driven approach reveals that the speed of listeners’ ability to categorize and properly label speech sounds is directly related to dynamic variations in their brain connectomics. These findings contribute in several ways to our understanding of how the brain works in categorical perception and provide a basis for further research. In future iterations of the work, we plan to improve our approach by including directional and dynamic connectivity analysis to better delineate the temporal emergence of the phenomena observed here.

## Acknowledgements

This work was supported by the National Institute on Deafness and Other Communication Disorders of the National Institutes of Health under award number NIH/NIDCD R01DC016267 (G.M.B.).

## APPENDIX

### GRAPH MINING

Mathematical definitions and interpretation of network features are given below:

#### Characteristics path

A fundamental property of brain networks is functional integration, which indicates how integrated a network is and, thus, how easily information flows (Rubinov and Sporns, 2010) among nodes. A widely-used approach to estimate properties of functional integration between nodes is based on the concept of characteristic path length. The characteristic path length is defined as the average shortest path length in the network (Watts and Strogatz, 1998). Hence, small characteristic path values imply dense connectivity and stronger potential for integration among nodes. Let, *L*_*i*_ is the average distance between node *i* and all other nodes of a network, Average Characteristic path is defined as:

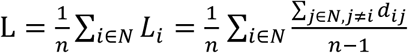

Where, *d*_*ij*_ is the shortest distance between node *i, j* (shortest path can be calculated using any popular shortest path algorithm), *N* is the set of all nodes, and *n* is the total number of nodes.

#### Global efficiency

Global efficiency (E) is used to find, how cost-efficient a particular network construction and how fault tolerant the network is. Hence, high global efficiency, implying the excellent use of resources. In brain connectivity analysis, structural and effective networks are similarly organized and share high global efficiency. On the other hand, functional networks have weaker connections and consequently share lower global efficiency (Honey et al., 2007). Global efficiency is the average of inverse shortest path length hence inversely related to the average characteristic path length. E is defined as:

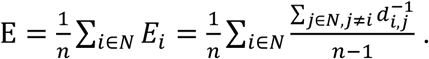

#### Average clustering coefficient

The average clustering coefficient for the network reflects, how close its neighbors are to being a clique or complete graph. The average clustering coefficient of a node is defined as the fraction of triangles around a node (Watts and Strogatz, 1998) and defined as:

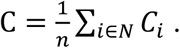

Here, C_i_ is the clustering coefficient of i^th^ node. Let k_i_ is the number of neighborhood node, and t_i_ is the number of triangles created around i^th^ node. If a node has k neighbors, there are *k(k* − 1)/2 edges could exist among the nodes within the neighborhood. Hence, C can be defined as:

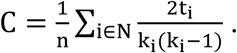

#### Transitivity

Transitivity is a classical variant of average clustering coefficient and defied as:

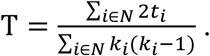

The value of average clustering coefficient can be influenced by nodes with a low degree. But transitivity is normalized collectively and consequently hence, does not have such problem (Newman, 2003).

#### Small-worldness

Small-world network (S) is formally defined as networks that are significantly densely clustered and have larger characteristic path length than random networks (Watts and Strogatz, 1998). Mathematically S can be expressed as:

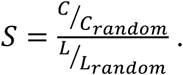

Where *C* and *C*_rand_ are the clustering coefficients, and *L* and *L*_rand_ are the characteristic path lengths of the test network and an equivalent random network with the same degree on average respectively. For a small world network S > 1, C >> C_random_ and L >> L_random_. Such network tends to contain more densely connected cliques/near-cliques/sub-networks than random network. Those sub-networks are interconnected by one or more edge.

#### Assortativity coefficient

Despite the importance local and community structure, it is essential to study global diversity in networks. Hence the tendency to connect nodes with similar numbers of edges. This tendency, called assortativity, described crucial dynamic and structural properties of real-world networks, such as epidemic spreading or error tolerance (Foster et al., 2010). A positive assortativity coefficient indicates that nodes tend to link to other nodes with the same or similar degree, on the other hand negative values indicate relationships between nodes of different degree. Biological networks typically show negative assortativity coefficient as high degree nodes tend to attach to low degree nodes (Piraveenan et al., 2012). Mathematically, the assortativity coefficient is the Pearson correlation coefficient of degree between pairs of linked nodes (Newman, 2002). Consider an undirected graph of N vertices and M edges with degree distribution *p*_*j*_. That is *p*_*j*_ is the probability that a randomly chosen node on the graph will have degree k and *q*_*k*_ is the distribution of the remaining degree. This *q*_*k*_ captures the number of edges leaving the node, other than the one that connects the pair. The assortativity coefficient (r) is defined as:

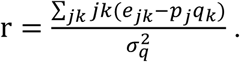

Where, 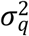 is the variance of distribution *p*_*k*_ and *e*_*jk*_ refers to the joint probability distribution of the remaining degrees of the two nodes.

#### Modularity Index

Modularity refers to the ability of subdivision the network into non-overlapping groups of nodes (known as modules or community) in a way that maximizes the number of within-group edges. Networks with high modularity have dense connections between the nodes within the modules but sparse connections between nodes in different modules. Hence, modularity quantifies the community strength of a test network by comparing the fraction of edges within the community with respect to random network (Chen et al., 2014). It is widely used to discover anatomical modules correspond to groups of specialized functional area which is previously determined by physiological recordings. Usually, anatomical, effective and functional modules in brain connectivity show extensive overlap (Rubinov and Sporns, 2010). Modularity index of a given network is the fraction of the edges that fall within the given groups minus the expected fraction if edges were distributed at random. Finding optimal modular structure is an optimization problem. Any optimization approach generally sacrifices some degree of accuracy for computational speed. Widely used algorithm to find optimal modular structure are proposed by Newman et al. (Newman, 2004), and Blondel et al. (Blondel et al., 2008).

#### Stability selection with Randomized Lasso

Randomized Lasso (RL) (Meinshausen and Bühlmann, 2010) is a straightforward two step approach. Instead of applying specific algorithm to the whole data set to determine the selected set of variables based on the weight of coefficient, RL applied randomized lasso several times to random subsamples of the data of size n/2 (n = number of samples) and chose those variables that are selected consistently across subsamples. By performing this double randomization several times, the method assigns high scores to features that are repeatedly selected across randomizations. In short, features selected more often are considered good features even though the “irrepresentable condition” (Zhao and Yu, 2006) is violated. This approach is similar to the concept of bagging (Breiman, 1999) and sub-bagging (Büchlmann and Yu, 2002) algorithm.

We know, Lasso has sparse solutions. For higher dimensional data, many estimated coefficients of variables become zero. Removing the variables can be used to reduce the dimensionality of the data. There are some limitations of Lasso-based feature selection are:

1. Lasso has a tendency to select an individual variable out of a group of highly correlated features.

2. When the correlation between features is not too high, the performance of Lasso is restrictive.

Lasso penalizes the absolute value of coefficients |*β*|_*k*_of every component with a penalty term proportional to the regularization parameter *λ* ∈ ℝ. On the other hand, Randomized Lasso penalizes using randomly chosen values in a range [*λ, λ*/*α*] where, *α* ∈ (0,1) is the weakness parameter. The concept of weakness parameter is closely related to weak greedy algorithms (Temlyakov, 2000). Let *W*_*k*_ be an i.i.d. random variable in a range from (*α*, 1) for k = 1, …., p. The estimator of Randomized Lasso can be written as (Meinshausen and Bühlmann, 2010):

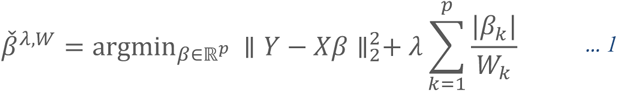

Here, Y and X is the class label and feature matrix respectively. Implementation of equation: 1 is a straightforward two-stage process:

1. Re-scaling of the feature variables (with scale factor W_k_ for the k-th variable),

2. LARS algorithm is applied on re-scaled variables (Efron et al., 2004).

In this approach, the reweighting is simply chosen at random. It is not sensible to expect improvement from randomization with one random perturbation. However, applying Randomized Lasso with many iterations (e.g. 1000 times) and looking for variables that are chosen frequently is a useful tool to find out stable feature (Meinshausen and Bühlmann, 2010).

By performing this double randomization several times, RL assigns high scores to features that are repeatedly selected across randomizations. if we run the Lasso for several bootstrapped replications of a given sample, then intersecting the supports of the Lasso bootstrap estimates leads to consistent model selection (Bach, 2008; Meinshausen and Bühlmann, 2010)

